# Ciliary phosphatidylinositol phosphatase Inpp5e plays positive and negative regulatory roles in Shh signaling

**DOI:** 10.1101/721399

**Authors:** Sandii Constable, Alyssa B. Long, Katharine A. Floyd, Stéphane Schurmans, Tamara Caspary

## Abstract

Sonic hedgehog (Shh) signal transduction specifies ventral cell fates in the neural tube and is mediated by the Gli transcription factors that play both activator (GliA) and repressor (GliR) roles. Cilia are essential for Shh signal transduction and the ciliary phosphatidylinositol phosphatase, Inpp5e, is linked to Shh regulation. In the course of a forward genetic screen for recessive mouse mutants, we identified a functional null allele of *Inpp5e*, *ridge top (rdg)*, with expanded ventral neural cell fates at E10.5. By E12.5, *Inpp5e*^*rdg/rdg*^ embryos displayed normal neural patterning and this correction over time required Gli3, the predominant repressor in neural patterning. *Inpp5e*^*rdg*^ function largely depended on the presence of cilia and on Smoothened, the obligate transducer of Shh signaling, indicating Inpp5e functions within the cilium to regulate the pathway. These data indicate that Inpp5e plays a more complicated role in Shh signaling than previously appreciated. We propose that Inpp5e attenuates Shh signaling in the neural tube through regulation of the relative timing of GliA and GliR production, which is important in understanding how duration of Shh signaling regulates neural tube patterning.

**Summary statement:** Inpp5e attenuates Sonic hedgehog signal transduction through a combination of positive and negative regulatory roles that likely control the relative timing of Gli processing.

## Introduction

Shh signaling plays a major role in determining the identity of the ventral cell fates in the developing neural tube (Echelard et al., 1993, Roelink et al., 1994). The cells at the ventral midline, called the floor plate, require a high concentration of Shh during a critical developmental time window (Ribes et al., 2010). Other ventral cell fates are specified due to both the concentration and duration of Shh signaling. For example, Nkx2.2-positive V3 interneuron precursors, can be specified by either high concentrations of Shh or by increasing time of exposure to lower amounts of Shh (Dessaud et al., 2007). The ability of a cell to monitor the duration of signaling is thus critical but we currently lack a detailed understanding of how cells interpret the duration of Shh signal.

The downstream effectors of Shh signaling are the Gli transcription factors Gli1, Gli2 and Gli3, which possess context-dependent activator and repressor functions. In the neural tube, Gli2 is the primary activator and Gli3 is the major repressor (Ding et al., 1998, Matise et al., 1998, Litingtung and Chiang, 2000, Persson et al., 2002). Full length Gli protein is processed to a mature activator (GliA) form or cleaved to a repressor (GliR) form based on the presence or absence of Shh ligand. GliR production in the absence of ligand depends on the orphan G-protein receptor Gpr161 increasing cAMP levels and protein kinase A (PKA) activity (Mukhopadhyay et al., 2013). PKA is crucial both to phosphorylate full length Gli at four specific sites which promotes its cleavage to GliR as well as to further phosphorylate full length Gli and prevent it from being processed to GliA (Mukhopadhyay et al., 2013, Niewiadomski et al., 2014). Thus, *Shh* mutants produce only GliR and do not specify ventral cell fates (Litingtung and Chiang, 2000, Chiang et al., 1996). In contrast, upon Shh stimulation, decreased ciliary cAMP levels antagonize PKA activity, preventing GliR and enabling GliA production (Moore et al., 2016). Thus, the physiological Shh response at any given position in the neural tube is determined by an effective Gli ratio, which integrates opposing gradients of activating GliA and repressive GliR.

The primary cilium is a microtubule-based projection found on almost all cells and is intimately associated with Shh signaling (Huangfu et al., 2003, Corbit et al., 2005, Rohatgi et al., 2007, Haycraft et al., 2005). Mutations in proteins required for cilia biogenesis and function lead to a breakdown in Shh signal transduction regulation and either an increase or an absence of ventral neural cell fates (reviewed in (Bangs and Anderson, 2017)). Several components of the Shh signaling pathway traffic to cilia and move dynamically upon pathway stimulation. These include the Shh receptor, Patched1 (Ptch1), and the obligate transducer of the pathway, Smoothened (Smo) (Corbit et al., 2005, Rohatgi et al., 2007). Smo enriches in cilia upon Shh stimulation and this enhanced localization is necessary but not sufficient for its activation. Concomitantly, Gli proteins are enriched at the ciliary tip (Haycraft et al., 2005). Upon genetic ablation of cilia, no ventral cell fates are specified because Gli proteins are not processed to either GliA or to GliR (Huangfu and Anderson, 2005, Liu et al., 2005).

The ciliary membrane is biochemically distinct from the plasma membrane as its phosphatidyl inositol (PI) composition is enriched for phosphatidyl inositol 4 phosphate (PI(4)P). (Chavez et al., 2015, Garcia-Gonzalo et al., 2015). PI(4)P is important in regulating the ciliary localization of Ptch1, as well as ciliary enrichment and activation of Smo (Jiang et al., 2016). In *Drosophila*, PI(4)P is converted between PI(4)P and PI by Sac1 phosphatase and Stt4 kinase. Sac1 mutants accumulate PI(4)P, leading to increased Shh signaling response (Yavari et al., 2010). Similarly, *ptc* mutants accumulate PI(4)P and increase Shh pathway activation (Yavari et al., 2010). This mechanism is conserved in mammalian cells where treatment of NIH3T3 cells with PI(4)P induces pathway activation along with Smo ciliary enrichment (Jiang et al., 2016). PI(4)P binds to both Ptch1 and Smo (Jiang et al., 2016). Stimulation with Shh increases binding of PI(4)P to Smo and decreases binding to Ptch1, making this molecule an attractive candidate to explain how Ptch1 inhibits Smo, a critical step in Shh signaling (Jiang et al., 2016).

The ciliary inositol polyphosphate-5-phosphatase E (Inpp5e) converts PI(3,4,5)P_3_ and PI(4,5)P_2_ (hereafter PIP_2_) to PI(3,4)P_2_ and PI(4)P respectively (Kisseleva et al., 2000). Inpp5e localizes to cilia where it maintains PI(4)P levels (Chavez et al., 2015, Garcia-Gonzalo et al., 2015, Jacoby et al., 2009, Bielas et al., 2009). Previous work investigated the role of Inpp5e in regulating Shh signaling. Mouse embryonic fibroblasts (MEFs) and neural stem cells lacking *Inpp5e* with either of two distinct null alleles, *Inpp5e*^*tm1.1Cmit*^ (hereafter *Inpp5e*^*ΔEx2-6*^) or *Inpp5e*^*tm1.2Ssch*^ (hereafter *Inpp5e*^*ΔEx7-8*^), display a diminished transcriptional response upon Shh stimulation (Chavez et al., 2015, Dyson et al., 2017, Garcia-Gonzalo et al., 2015). This correlates with increased ciliary PIP_2_ levels recruiting Tulp3 and the Shh antagonist Gpr161 into cilia (Garcia-Gonzalo et al., 2015, Dyson et al., 2017). Gpr161 resides in the cilium in the absence of Shh and generates cAMP (Mukhopadhyay et al., 2013). Increased levels of cAMP activate PKA which phosphorylates full length Gli2 and Gli3 resulting in cleavage to GliR and repression of the Shh response (Tuson et al., 2011, Wang et al., 2000). Consistent with this, the ventral-most neural cell fates requiring the highest level of Shh response are lost in *Inpp5e*^*ΔEx2-6/ΔEx2-6*^ embryos (Dyson et al., 2017). The simplest interpretation of these data is that Inpp5e normally plays an activating role in Shh signaling. However, ventral neural cell fates requiring intermediate levels of Shh response are dorsally expanded in the *Inpp5e*^*ΔEx2-6/ΔEx2-6*^ mutants suggesting a more complicated mechanism is at play (Dyson et al., 2017).

Here, we investigate the role of Inpp5e in the Shh signaling pathway using an *Inpp5e* point mutant we identified in a forward genetic screen. We genetically demonstrate that this mutation generates a functional null allele and that loss of Inpp5e activity leads to an initial expansion of ventral neural tube cell fates indicating Inpp5e negatively regulates the Shh response. Interestingly, we found that the ventral pattern corrects over time and we demonstrate this mechanism requires Gli3, the predominant repressor in neural patterning. Inpp5e localizes to cilia, which we show are required for Inpp5e function in regulating Shh signaling. We found that when Inpp5e function is absent, Smo is required for the highest Shh response but not for more intermediate Shh patterning responses. From these data, we propose that Inpp5e plays a critical role in controlling the level of Shh response over time by attenuating the pathway through control of the timing of GliA/GliR gradient production. These data provide a mechanistic framework from which to understand how the duration of Shh signaling regulates ventral neural cell fate.

## Results

### Inpp5e negatively regulates the Shh signaling response in the E10.5 neural tube

We previously published the *Inpp5e*^*M2*^ line, and showed it carried an A-to-G transition in the mouse *Inpp5e* gene as well as a change in the *Slc2a6* gene 0.625 Mb away (Sun et al., 2012). To eliminate the *Slc2a6* mutation, we backcrossed to FVB and generated the recombinant chromosome carrying only the *Inpp5e* mutation. To distinguish this recombinant line from the *Inpp5e*^*M2*^ line, we refer to this line as *ridge top (rdg)*, *Inpp5e*^*rdg*^, to reflect the exencephaly that resembles a ridgetop hat.

We evaluated neural tube patterning in *Inpp5e*^*rdg/rdg*^ mutant embryos at embryonic day (E) 10.5 by staining sections with antibodies for specific cell fates. Shh is initially expressed in the notochord and subsequently in the floor plate, which also expresses FoxA2 (Echelard et al., 1993, Roelink et al., 1994, Sasaki and Hogan, 1994, Riddle et al., 1993). In wild type and *Inpp5e*^*rdg/rdg*^ mutants, the floor plate was visible and expressed both Shh and FoxA2 (Fig. 1A, B, F, G). Additionally, we observed FoxA2 positive cells that did not co-express Shh scattered dorsally within the ventricular zone in *Inpp5e*^*rdg/rdg*^ mutants (Fig. 1G). In wild type sections, p3 cells adjacent to the floorplate expressed Nkx2.2 whereas in *Inpp5e*^*rdg/rdg*^ mutant sections, Nkx2.2-positive cells expanded dorsally in a dispersed manner, similar to what we observed with FoxA2 (Fig. 1C, H). Additional ventral cell fates expressing Nkx6.1 and Olig2 also expanded dorsally in the *Inpp5e*^*rdg/rdg*^ mutants (Fig. 1H, I). The dorsal boundary for all the expanded cell fates correlated to the ventral boundary of Pax6, which is repressed by Shh signaling (Lek et al., 2010, Ericson et al., 1997). This boundary is shifted dorsally in *Inpp5e*^*rdg/rdg*^ mutants compared to wild type controls (Fig. E, J). Taken together, these data demonstrate an expansion of Shh-dependent ventral cell fates in *Inpp5e*^*rdgrdg*^ mutant embryos, suggesting an expansion of the Shh activity gradient in the *Inpp5e*^*rdg/rdg*^ neural tube.

**Figure 1:**
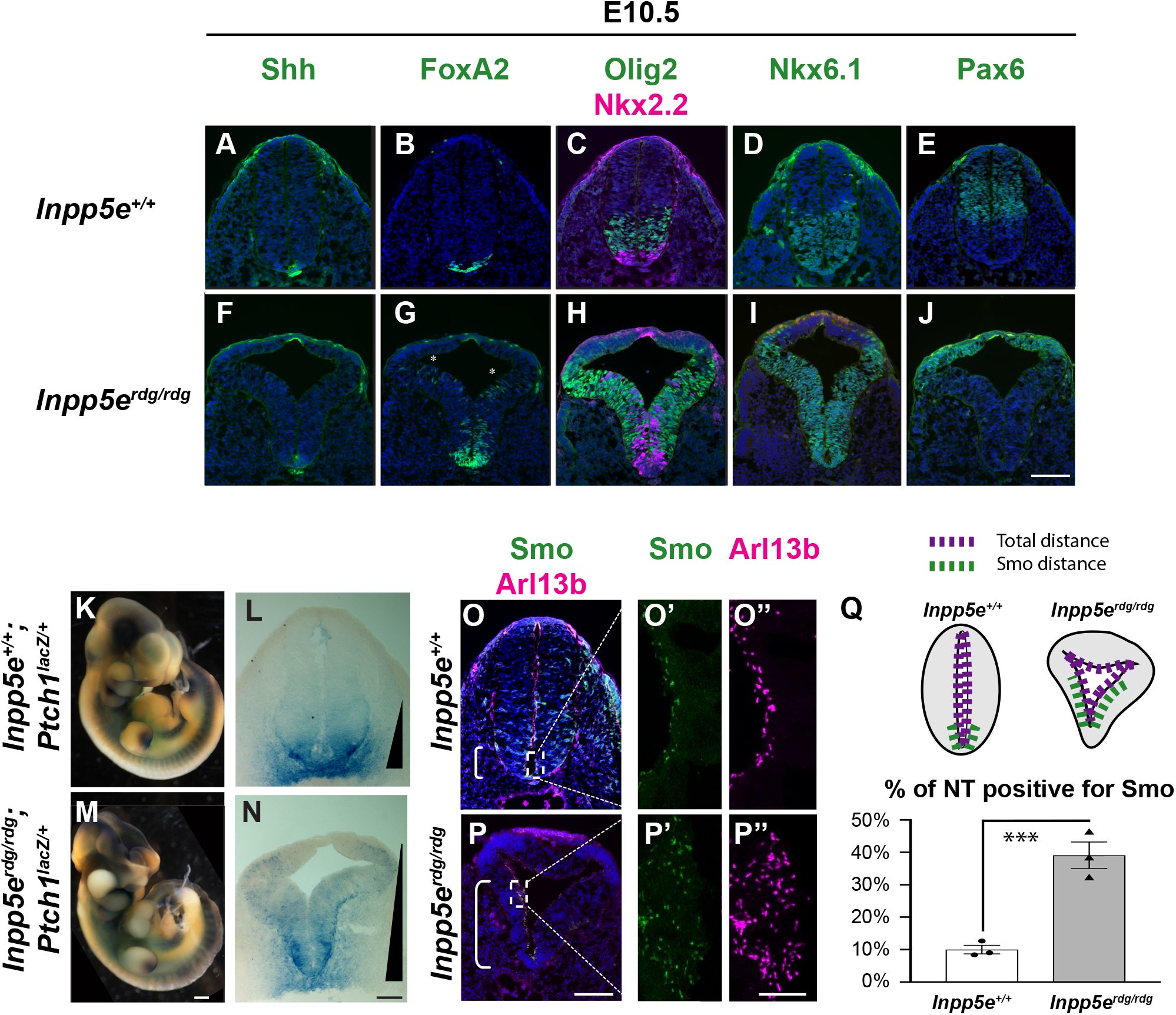
Inpp5e negatively regulates the Shh signaling response in the E10.5 neural tube. (A-J) Caudal (hindlimb) sections of E10.5 wild type (A-E, n=4) and *Inpp5e*^*rdg/rdg*^ (F-J, n=4) neural tubes stained with antibodies against the indicated transcription factors. (A, F) *Inpp5e*^*rdg/rdg*^ mutants display normal Shh expression in the floor plate. (B, G, C, H, D, I) FoxA2, Olig2, Nkx2.2 and Nkx6.1 are expanded dorsally in *Inpp5e*^*rdg/rdg*^ mutants. The FoxA2, Olig2 and Nkx2.2 domains are intermixed with a few dorsally scattered FoxA2 cells (G, asterisks). (E, J) The Pax6 domain is shifted dorsally in *Inpp5e*^*rdg/rdg*^ mutants compared to wild type littermates. (K, M) Whole mount and (L, N) neural tube sections of *Inpp5e*^+/+^;*Ptch1*^*LacZ/+*^ (n=5) and *Inpp5e*^*rdg/rdg*^;*Ptch1*^*LacZ/+*^ embryos (n=5) stained for β-galactosidase activity, which reflects Shh activity. (K-N) *Inpp5e*^*rdg/rdg*^;*Ptch1*^*LacZ/+*^ mutants display a dorsal expansion of β-galactosidase activity in the neural tube compared to wild type littermates. Extent of gradient indicated by black triangle. (O-P) Wild type (n=3) and *Inpp5e*^*rdg/rdg*^ (n=3) neural tube sections stained with antibodies against Smo and cilia marker Arl13b. Bracket depicts region containing Smo-positive cilia. (O’, O’’, P’ P’’) Enlarged view of cilia from region indicated by dotted boxes in O and P. (Q) Schematic describing quantification of region of Smo staining and graph reflecting percentage of ventricular lumen displaying ciliary Smo enrichment (see Methods for details). Values displayed are mean ± SEM of three biological replicates analyzed by two tailed unpaired t-test with Welch’s correction. ***p<0.001. Scale bars: 100μm for A-J, L, N, O, and P; 400μm for K and M; 10μm for O’, O’’, P’, and P”.

In order to monitor the Shh activity gradient, we used a *Ptch1*^*LacZ*^ allele as *Ptch1* is a transcriptional target of the Shh signaling pathway (Goodrich et al., 1997, Goodrich et al., 1996). Upon X-gal staining of whole E10.5 wild type and *Inpp5e*^*rdg/rdg*^ mutant embryos, we detected blue staining reflecting the Shh transcriptional response (Fig. 1K, M). In wild type embryos, we observed staining in known Shh signaling centers including the notochord and the zone of polarizing activity in the limb buds (Fig. 1K). These regions also stained blue in the *Inpp5e*^*rdg/rdg*^; *Ptch1*^*LacZ/+*^ mutant embryos indicating Shh activity (Fig. 1M). In neural tube sections of wild type *Ptch1*^*LacZ/+*^ embryos, we saw a steep *LacZ* expression gradient with the strongest blue staining at the ventral midline of the neural tube (Fig. 1L). In contrast, the staining expanded dorsally in the *Inpp5e*^*rdg/rdg*^; *Ptch1*^*LacZ/+*^ mutant embryos (Fig. 1N), consistent with the dorsal expansion of ventral cell fates we observed and indicating dorsally expanded Shh activity in *Inpp5e*^*rdg/rdg*^; *Ptch1*^*LacZ/+*^ mutants.

At a mechanistic level, Smo ciliary enrichment correlates with Smo activation, so we examined Smo staining in neural tube cilia (Corbit et al., 2005). In wild type sections, we observed ciliary Smo enrichment near the ventral midline (Fig. 1O). However, in *Inpp5e*^*rdg/rdg*^ embryos, we found Smo-positive cilia expanded dorsally along the ventricular zone (Fig. 1P). The *Inpp5e*^*rdg/rdg*^ neural tube shape is abnormal, almost triangular. Thus, to quantify the Smo expansion, we drew a line along the entire neural tube lumen to calculate total luminal distance. We found no statistical difference in total neural tube distance between wild type and *Inpp5e*^*rdg/rdg*^ suggesting the *Inpp5e*^*rdg/rdg*^ neural tube is simply misshapen (p>0.5). We then measured the distance over which we observed ciliary Smo enrichment and report this as a percentage of the total distance of the neural tube lumen (Fig. 1Q). We found ciliary Smo enrichment in the ventral 10% of the neural tube in wild type sections and in the ventral 40% of the neural tube in *Inpp5e*^*rdg/rdg*^ sections (p<0.001) (Fig. 1Q). This dorsal expansion of ciliary Smo enrichment correlates with both the expanded Shh response shown with the *Ptch1-LacZ* reporter and the dorsal expansion of cell fates shown by immunofluorescence, suggesting that the ciliary Smo is activated. Taken together, these data indicate that the normal role of Inpp5e is to negatively regulate the Shh signaling response in the E10.5 neural tube.

### Inpp5e^rdg^ is a functional null allele sensitive to strain background

The *Inpp5e*^*rdg*^ allele changed aspartic acid to glycine at residue 511 within the phosphatase domain of Inpp5e, Inpp5e^D511G^ (Fig. 2A, B). This region of the protein sequence is highly conserved across species (Fig. 2C). At E12.5, most *Inpp5e*^*rdg/rdg*^ embryos exhibited exencephaly (88%, 61/69) and either microphthalmia or anophthalmia (80%, 33/41). 77% (31/40) of embryos exhibited a rough neural tube with 30% showing spina bifida at the hindlimb level (13/40) (Fig. 2E). The *Inpp5e*^*rdg/rdg*^ embryos died around E13.5. In order to determine whether the D511G mutation was causative, we crossed *Inpp5e*^*rdg/+*^ to mice carrying an *Inpp5e* deleted allele (*Inpp5e*^*ΔEx7-8/+*^) and examined the compound heterozygote embryos. *Inpp5e*^*rdg/ΔEx7-*8^ embryos display exencephaly, anophthalmia, spina bifida and a rough neural tube indicating the alleles failed to complement and that *Inpp5e*^*rdg*^ is an allele of *Inpp5e* (Fig. 2F).

**Figure 2:**
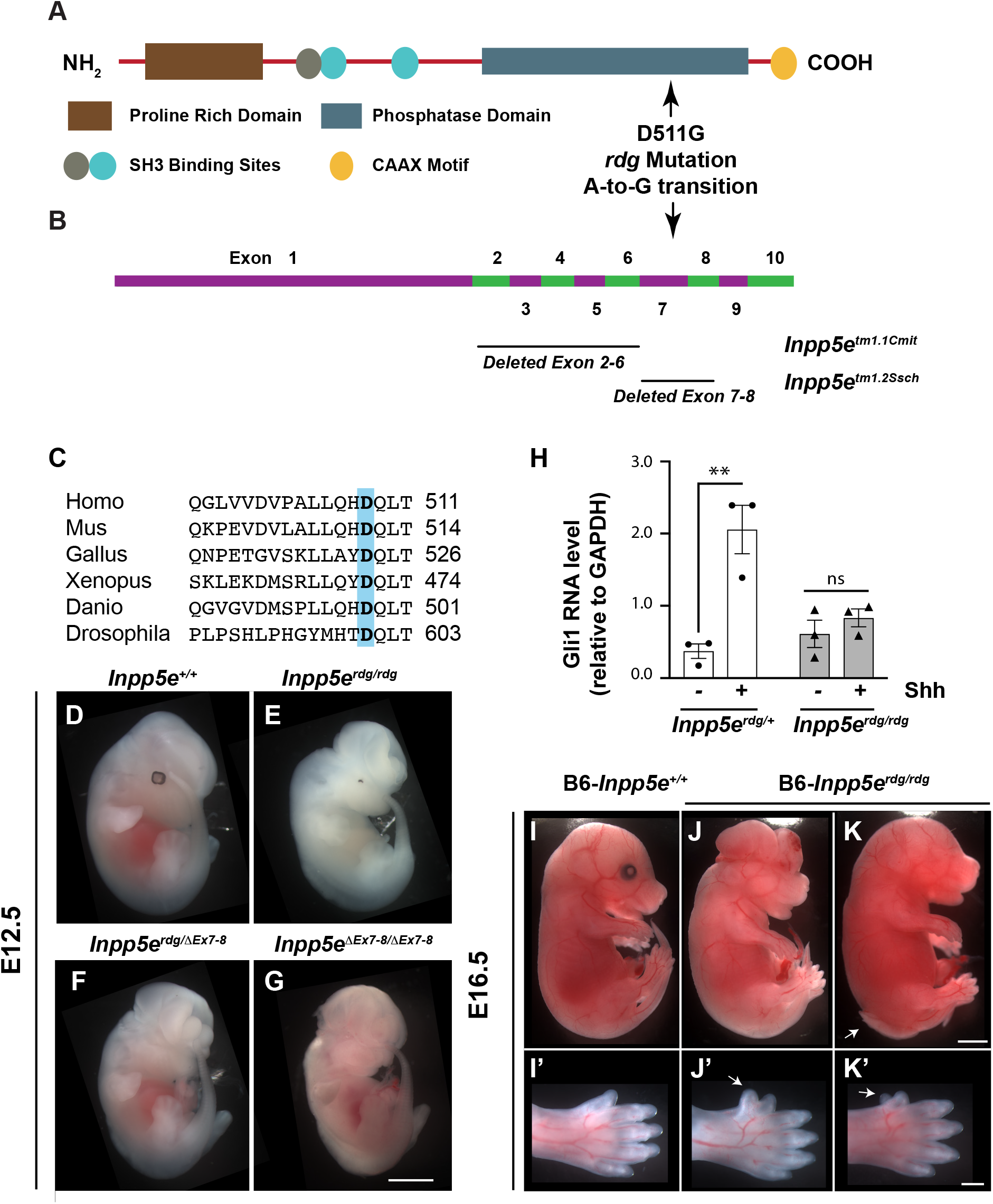
*Inpp5e*^*rdg*^, carrying a D511G mutation, is a functional null allele of *Inpp5e*. (A) Schematic showing the protein domain structure of Inpp5e, including the location of the amino acid change. *Inpp5e*^*rdg*^ contains an aspartic acid to glycine change at position 511 as a result of an A-to-G transition in the coding strand (NM_033134.3:c2164A>G). (B) Exon structure of *Inpp5e*, based on NCBI reference sequence NM_033134.3, showing location of deleted exons in two previously characterized alleles. Note: diagrams in A and B are aligned, indicating that D511G is located within exon 7, in Inpp5e’s phosphatase domain. (C) Alignment of protein sequence surrounding position 511 of mouse Inpp5e showing the aspartic acid is conserved across multiple species. (D-G) Whole mount images of E12.5 embryos. (E) *Inpp5e*^*rdg/rdg*^ mutants display exencephaly and microphthalmia (n=20). Note: these animals are on a mixed FVB/C3H background so that the retinal pigment epithelium cells are visible. (F) *Inpp5e*^*rdg/ΔEx7-8*^ embryos resemble *Inpp5e*^*rdg/rdg*^ mutants, indicating the alleles fail to complement (n= 4). (G) *Inpp5eΔEx7-8/ΔEx7-8* mutants have the same appearance as *Inpp5e*^*rdg/rdg*^ (n=9). (H) qRT-PCR of *Gli1*, a Shh transcriptional target, in *Inpp5e*^*rdg/+*^ and *Inpp5e*^*rdg/rdg*^ MEFs in the presence (+) and absence (-) of Shh treatment. *Inpp5e*^*rdg/rdg*^ MEFs do not respond to Shh. Values displayed are mean ± SEM, analyzed by two-way ANOVA using Tukey correction for multiple comparisons. **p<0.01. ns, not significant. Data is from three independent experiments. (I-K) *Inpp5e*^*rdg/rdg*^ mutants on C57BL/6J background survive to E16.5 and display exencephaly, microphthalmia, and spina bifida to varying degrees of severity (n=6). Arrow points to spina bifida in caudal neural tube. (I’-K’) Close up images showing hindlimb preaxial polydactyly. Arrow indicates extra digit. Scale bars: whole embryo 1mm, hindlimb 500μm.

Upon stimulation of cultured cells with the Shh agonist SAG, cells lacking Inpp5e show a diminished Shh response (Chavez et al., 2015, Dyson et al., 2017, Garcia-Gonzalo et al., 2015). To determine the Shh response of cells carrying *Inpp5e*^*rdg*^, we generated MEFs from *Inpp5e*^*rdg/rdg*^ and control littermate embryos and tested their ability to respond to Shh stimulation. As expected, we found control MEFs increased expression of *Gli1*, a Shh target gene, upon stimulation with Shh-conditioned media (Fig. 2H) (p<0.01). However, the *Inpp5e*^*rdg/rdg*^ MEFs displayed no change in *Gli1* levels when treated with Shh (Fig. 2H). Similar results have been shown for cell lines lacking Inpp5e function (Chavez et al., 2015, Dyson et al., 2017, Garcia-Gonzalo et al., 2015). Thus, the *Inpp5e*^*rdg*^ allele phenocopies *Inpp5e* null alleles in cultured cells.

The fact that *Inpp5e*^*rdg/rdg*^ embryos died at E13.5 contrasts with the two previously studied *Inpp5e* null alleles: *Inpp5e*^*ΔEx7-8*^ which deletes exons 7-8 and *Inpp5e*^*ΔEx2-6*^ which deletes exons 2-6 (Fig. 2B). Both of these deletion mutants die at birth and both were analyzed on a predominantly C57BL/6 background (with some possible contribution from 129/Sv) (Dyson et al., 2017, Jacoby et al., 2009). Our analysis of the *Inpp5e*^*rdg*^ allele was on an FVB background. To distinguish whether such phenotypic distinctions reflected differences in allelic function or strain background, we backcrossed the *Inpp5e*^*ΔEx7-8*^ allele onto FVB for three generations. We found that these FVB-*Inpp5e*^*ΔEx7-8/ΔEx7-8*^ embryos died at E13.5-14.5 and displayed exencephaly, anophthalmia, spina bifida and a rough neural tube (Fig. 2G). We also examined neural tube patterning in E10.5 FVB-*Inpp5e*^*ΔEx7-8/ΔEx7-8*^ embryos and found expanded ventral neural cell fates comparable to *Inpp5e*^*rdg/rdg*^ (Fig. S1A-J). Thus, the *Inpp5e*^*ΔEx7-8*^ allele on the FVB background phenocopies the *Inpp5e*^*rdg*^ allele. We performed the reciprocal experiment and crossed the *Inpp5e*^*rdg*^ allele onto the C57BL/6 background for four generations. We identified live B6-*Inpp5e*^*rdg/rdg*^ embryos at E16.5 (6/25, 24%) that displayed exencephaly (5/6), spina bifida (2/6), microphthalmia or anophthalmia (6/6) and hindlimb preaxial polydactyly (5/6) (Fig. 2I-K’). In contrast, we found no *Inpp5e*^*rdg/rdg*^ embryos at E16.5 on the FVB background (0/51). Taken together, these data indicate that Inpp5e-dependent phenotypes are sensitive to genetic background and that *Inpp5e*^*rdg*^ is a functional null allele.

### Normal ventral neural patterning in E12.5 Inpp5e^rdg/rdg^ mutant embryos indicates recovery of patterning over time

In the course of comparing the phenotypes of the *Inpp5e*^*rdg*^ and *Inpp5e*^*ΔEx7-8*^ alleles, we evaluated neural patterning at E12.5. In *Inpp5e*^*rdg/rdg*^ embryos, we found FoxA2 expression was restricted to the floor plate as in control embryos (Fig. 3A, E). Similarly, we found Nkx2.2-positive cells only in the p3 domain adjacent to the floor plate in control and *Inpp5e*^*rdg/rdg*^ mutant embryos (Fig. 3B, F). Overall, Olig2- and Nkx6.1-positive cells appeared in their normal domains with only slight expansion of the pMN domain of *Inpp5e*^*rdg/rdg*^ embryos compared to controls (Fig. 3B, C, F, G). These data indicate that the Shh response in *Inpp5e*^*rdg/rdg*^ mutant embryos is comparable to wild type by E12.5. We also observed normal patterning in FVB-*Inpp5e*^*ΔEx7-8/ΔEx7-8*^ embryos, indicating the recovery of the Shh response also occurs in this allele (Fig. S1K-T).

**Figure 3:**
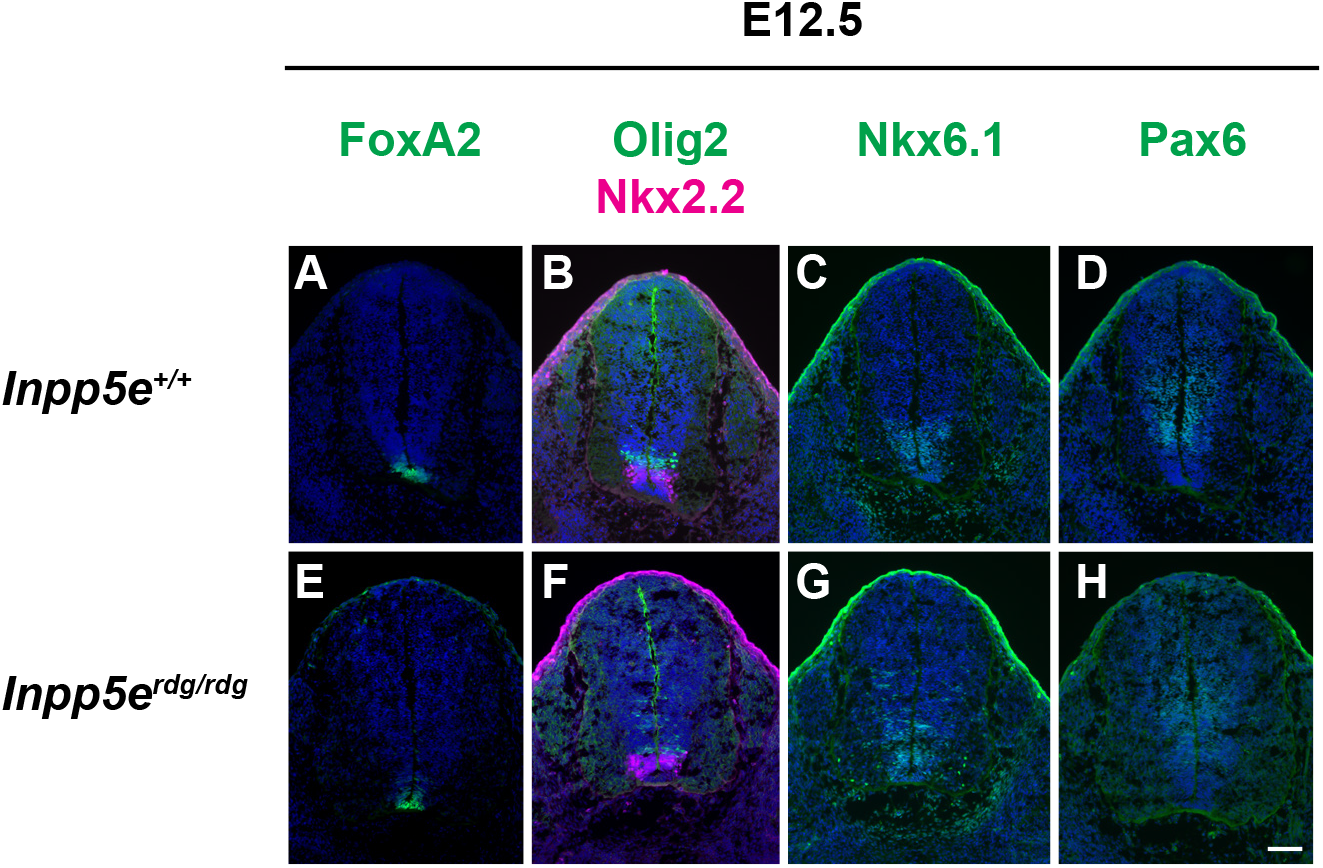
Normal ventral neural patterning in E12.5 *Inpp5e*^*rdg/rdg*^ embryos reflects a recovery of Shh response. Caudal (hindlimb) sections of E12.5 wild type (A-D) and *Inpp5e*^*rdg/rdg*^ (E-H) neural tubes stained with antibodies against the indicated cell fates. (A-B, D, E-F, H) FoxA2, Nkx2.2 and Pax6 expression domains look similar in wild type and *Inpp5e*^*rdg/rdg*^ neural tube sections (wild type n=6, *Inpp5e*^*rdg/rdg*^ n=5). (B-C, F-G) The Olig2 and Nkx6.1 domains in *Inpp5e*^*rdg/rdg*^ neural tubes show only a few cells scattered dorsally compared to wild type littermates. Scale bar: 100μm.

Pax6 expression is repressed by Shh signaling (Ericson et al., 1997). In order to test whether the dorsal boundary of the Shh response is normal in *Inpp5e*^*rdg/rdg*^ and *Inpp5e*^*ΔEx7-8/ΔEx7-8*^ embryos, we stained neural tube sections with antibody against Pax6. We found Pax6 expression in *Inpp5e*^*rdg/rdg*^ and *Inpp5e*^*ΔEx7-8/ΔEx7-8*^ embryos appeared the same as in wild type controls (Fig. 3D, H; Fig. S1K-T). Thus, the abnormal ventral patterning we observed in E10.5 *Inpp5e*^*rdg/rdg*^ and *Inpp5e*^*ΔEx7-8/ΔEx7-8*^ embryos recovers by E12.5. Taken together, these data imply that Inpp5e is not simply a negative regulator of the Shh response in the neural tube.

### Altered ciliary enrichment of Tulp3 and Gpr161 in Inpp5e^rdg/rdg^ neural tubes

Inpp5e removes the 5-phosphate from PI(3,4,5)P_3_ and PI(4,5)P_2_ and loss of Inpp5e function results in increased ciliary PIP_2_ (Garcia-Gonzalo et al., 2015, Chavez et al., 2015, Kisseleva et al., 2000). Tulp3 traffics G-protein coupled receptors (GPCRs) into cilia in a PIP_2_-dependent manner (Mukhopadhyay et al., 2010). Increased PIP_2_ levels in Inpp5e-deficient cilia increases the amount of Tulp3 seen (Chavez et al., 2015, Garcia-Gonzalo et al., 2015). To determine whether *Inpp5e*^*rdg*^ affected Tulp3 localization in a similar manner, we stained E10.5 neural tube sections with antibodies against Tulp3. We detected no Tulp3 in wild type neural tube cilia (Fig. 4A-C). In contrast, we observed Tulp3 staining in 90% of Arl13b positive luminal cilia in the ventral neural tube of the *Inpp5e*^*rdg/rdg*^ mutants (Fig. 4D-G). As Tulp3 is a known PIP_2_-binding protein, this result is consistent with *Inpp5e*^*rdg*^ being a loss-of-function allele that increases ciliary PIP_2_.

**Figure 4:**
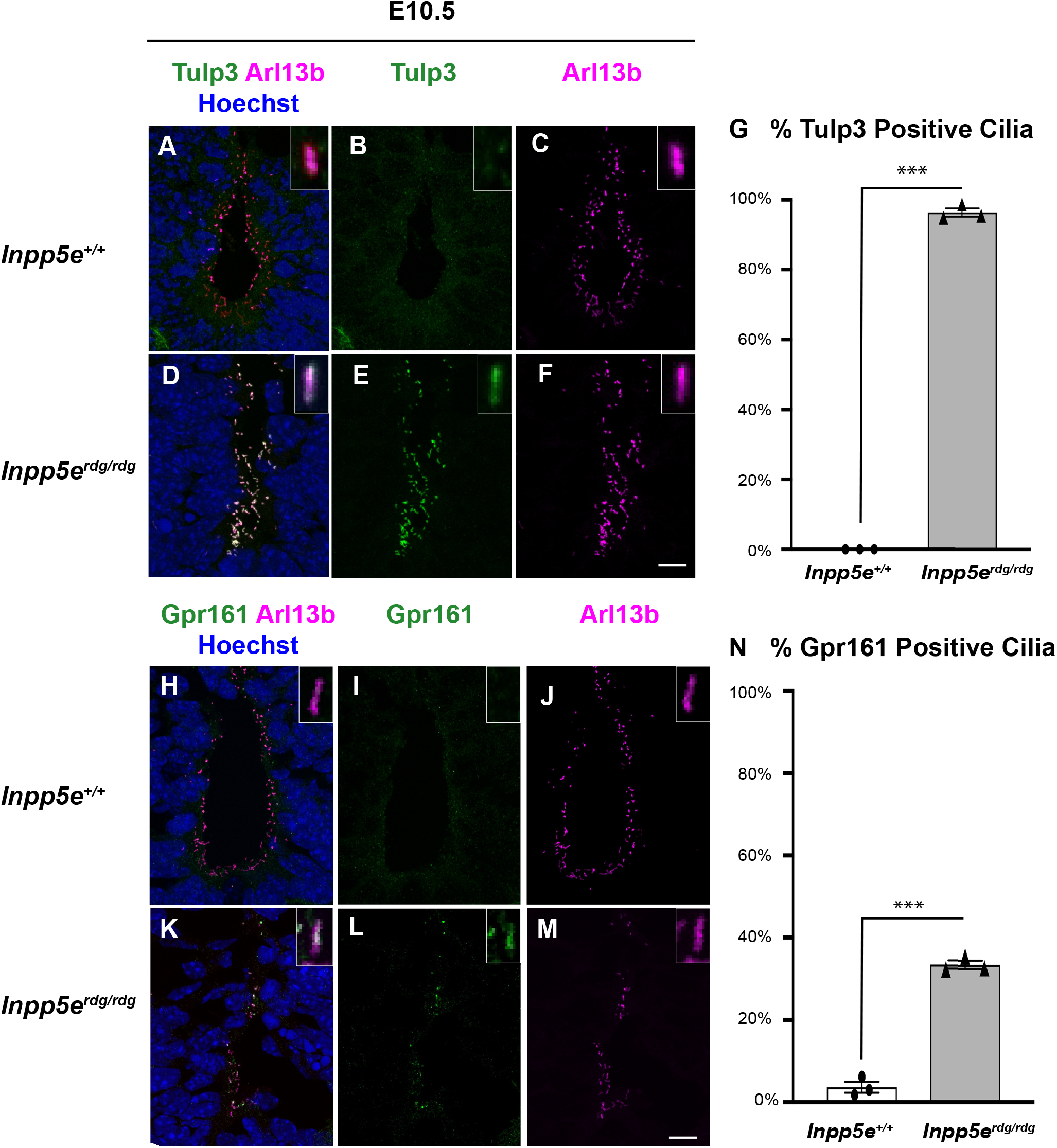
Altered ciliary Tulp3 and Gpr161 in *Inpp5e*^*rdg/rdg*^ neural tubes. (A-F, H-M) Images of cilia found in the ventral most region of E10.5 caudal (hindlimb) sections of wild type (A-C, H-J) and *Inpp5e*^*rdg/rdg*^ (D-F, K-M) neural tubes stained with antibodies against Tulp3, Gpr161 and Arl13b. Insets show a single cilium digitally magnified. (A-F) Tulp3 is found in almost all cilia in the neural tube of *Inpp5e*^*rdg/rdg*^ embryos but is absent from wild type cilia (wild type n=3, *Inpp5e*^*rdg/rdg*^ n=3). (G) Graphical representation of percentage of Arl13b-positive cilia also staining for Tulp3 (see Methods for details). (H-M) Gpr161 is found in an increased number of cilia in the neural tube of *Inpp5e*^*rdg/rdg*^ when compared to wild type (wild type n=3, *Inpp5e*^*rdg/rdg*^ n=3). (N) Quantification of the percentage of Arl13b-positive cilia also staining for Gpr161 (see Methods for details). Values displayed are mean ± SEM of three biological replicates analyzed by two tailed unpaired t-test with Welch’s correction. *** p<0.001. Scale bar: 10μm.

Increased PIP_2_ in Inpp5e-deficient cilia recruits Shh antagonist Gpr161 via a PIP_2_/Tulp3-dependent mechanism (Chavez et al., 2015, Garcia-Gonzalo et al., 2015). To determine whether an increase in ciliary levels of Gpr161 occurred coincident with the increase in Tulp3, we examined the localization of Gpr161 in ventral neural tube cilia. In wild type embryos, 3.4% of cilia were Gpr161-positive, whereas 34% of cilia stained for Gpr161 in *Inpp5e*^*rdg/rdg*^ embryos (p<0.001) (Fig. 4H-N). These data imply that in the neural tube, the increased PIP_2_ in *Inpp5e*^*rdg/rdg*^ mutant cilia efficiently recruits Tulp3 to cilia but Tulp3 is not sufficient to recruit Gpr161 to all cilia *in vivo*.

### Ift172-dependent and -independent functions of Inpp5e

Inpp5e is predominately localized to the cilium (Bielas et al., 2009, Jacoby et al., 2009), and Shh signaling is tightly associated with the cilium (Huangfu et al., 2003, Goetz and Anderson, 2010). To determine whether the *Inpp5e*^*rdg/rdg*^ neural tube patterning phenotype requires cilia, we generated *Inpp5e*^*rdg/rdg*^;*Ift172*^*wim/wim*^ double mutant embryos. Ift172 is an intraflagellar transport protein required for ciliary assembly and maintenance. *Ift172*^*wim/wim*^ single mutants do not produce a cilium, show exencephaly without a groove at the ventral midline and do not specify ventral neural cell fates, with the exception of Nkx6.1-positive cells which are Shh-dependent and cilia-independent (Fig. 5E,F) (Huangfu et al., 2003, Norman et al., 2009, Briscoe et al., 2000). In contrast, *Inpp5e*^*rdg/rdg*^ single mutants displayed pronounced exencephaly with a prominent groove at the midline (Fig. 5C). In the neural tube of *Inpp5e*^*rdg/rdg*^ mutants at E9.5, we observed FoxA2 expression at the ventral midline which expanded dorsally albeit diffusely through the majority of the neural tube (Fig. 5D). We identified an additional dorsal expansion of ventral cells expressing Nkx2.2 and Olig2 intermingled with the FoxA2-positive cells (Fig. 5D, D’). We found *Inpp5e*^*rdg/rdg*^;*Ift172*^*wim/wim*^ double mutant embryos resemble *Ift172*^*wim/wim*^ embryos showing exencephaly lacking a ventral midline groove (Fig. 5G). In the neural tube, we found *Inpp5e*^*rdg/rdg*^;*Ift172*^*wim/wim*^ double mutants specified no FoxA2-or Nkx2.2-positive cells, and had normal numbers of Nkx6.1-positive cells similar to *Ift172*^*wim/wim*^ single mutants (Fig. 5H, H’, H’’), suggesting that the *Inpp5e*^*rdg/rdg*^ phenotype is Ift172-dependent and that Inpp5e functions within cilia. However, we observed Olig2-positive cells in *Inpp5e*^*rdg/rdg*^;*Ift172*^*wim/wim*^ double mutants similar to the pattern in *Inpp5e*^*rdg/rdg*^ single mutants, suggesting some Ift172-independent Inpp5e function.

**Figure 5:**
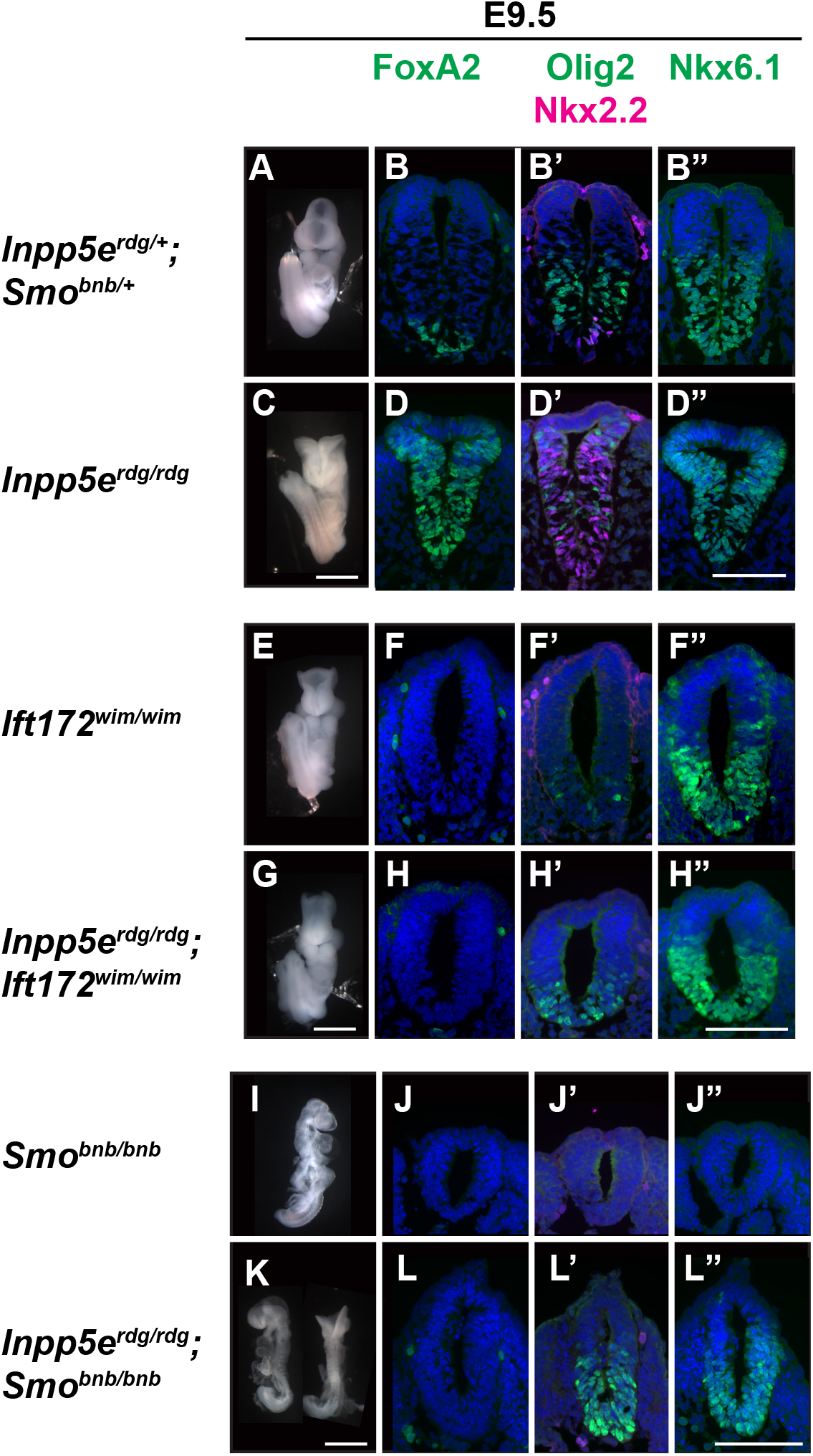
Inpp5e is cilia dependent and depends on Smoothened to specify cell fates requiring the highest Shh activity, but not those requiring moderate Shh activity. E9.5 whole mount images and neural tube staining of caudal (hindlimb) sections with indicated antibodies. (A, C) Ventral view of control and *Inpp5e*^*rdg/rdg*^ embryos highlighting exencephaly with prominent midline groove in the mutant (*Inpp5e*^*rdg/rdg*^ n=63). (B-B’’, D-D’’) Neural tube staining of control and *Inpp5e*^*rdg/rdg*^ embryos show ventral expansion of FoxA2, Nkx2.2, Olig2 and Nkx6.1 cell fates in mutant (wild type n=4, *Inpp5e*^*rdg/rdg*^ n=4). (E) Ventral view of *Ift172*^*wim/wim*^ embryo showing exencephaly without midline groove (n=20). (G) *Inpp5e*^*rdg/rdg*^;*Ift172*^*wim/wim*^ mutants display exencephaly that resembles the *Ift172*^*wim/wim*^ mutant (n=6). (F-F’’, H-H’’) Neural tube staining of both *Ift172*^*wim/wim*^ and *Inpp5e*^*rdg/rdg*^;*Ift172*^*wim/wim*^ show that these mutants fail to specify ventral cell fates, indicating the *Inpp5e*^*rdg/rdg*^ phenotype is cilia dependent (*Ift172*^*wim/wim*^ n=1, *Inpp5e*^*rdg/rdg*^;*Ift172*^*wim/wim*^ n=3). (I) Side view of *Smo*^*bnb/bnb*^ mutants reveals a small unturned body with a closed head (n=73). (K) Side and dorsal views of *Inpp5e*^*rdg/rdg*^;*Smo*^*bnb/bnb*^ embryos showing unturned bodies similar to *Smo*^*bnb/bnb*^ (n=9) although some have exencephaly, as seen in the dorsal view (n=5/9). (J-J’’) Neural tube staining confirms *Smo*^*bnb/bnb*^ embryos do not specify ventral cell fates as they cannot transduce the Shh response (n=1). (L-L’’) *Inpp5e*^*rdg/rdg*^;*Smo*^*bnb/bnb*^ mutants lack floor plate (FoxA2) and p3 (Nkx2.2) specification, indicating Inpp5e requires Smo to specify cell fates requiring the highest levels of Shh activity. In contrast, *Inpp5e*^*rdg/rdg*^;*Smo*^*bnb/bnb*^ mutants specify Olig2 and Nkx6.1 cells, indicating Inpp5e functions independently of Smo to specify cell fates requiring moderate Shh activity (n=5). Scale bars: whole embryo 1mm, neural tube sections 100μm.

### Smo-dependent and -independent functions of Inpp5e

Ciliary Smo enrichment correlates with Smo activation (Corbit et al., 2005). Given that Inpp5e appeared to act within the cilium, we explored the relationship of Inpp5e and Smo. To do this, we intercrossed *Inpp5e*^*rdg/+*^;*Smo*^*bnb/+*^ transheterozygous animals which should generate double mutant embryos at a frequency of 1 out of 16 embryos. We identified fewer *Inpp5e*^*rdg/rdg*^;*Smo*^*bnb/bnb*^ mutants (3%) than expected (6.25%), which is statistically significant (Chi-squared test, p= 0.02). Smo *bentbody*, *Smo*^*bnb*^, is a null allele and *Smo*^*bnb/bnb*^ embryos have a closed, misshapen head and fail to complete embryonic turning before dying by E9.5 (Caspary et al., 2002) (Fig. 5I). In neural tube sections, *Smo*^*bnb/bnb*^ lack ventral cell fate specification so do not express the Shh-dependent cell markers FoxA2, Nkx2.2, and Olig2 (Fig. 5J, J’). At E9.5, *Inpp5e*^*rdg/rdg*^;*Smo*^*bnb/bnb*^ double mutant embryos were small with partially turned or unturned bodies similar to *Smo*^*bnb/bnb*^ mutants (66%, 6/9). Unlike *Smo*^*bnb/bnb*^ single mutant embryos, *Inpp5e*^*rdg/rdg*^;*Smo*^*bnb/bnb*^ double mutant embryos displayed exencephaly (55%, 5/9) (Fig. 5K). In the neural tube of *Inpp5e*^*rdg/rdg*^;*Smo*^*bnb/bnb*^ mutants, we detected no expression of FoxA2 or Nkx2.2, cell fates requiring the highest Shh response (Fig. 5L, L’). The lack of ventral cell fates resembled the *Smo*^*bnb/bnb*^ mutant and indicated that Inpp5e requires Smo function to specify these cell fates. In contrast, unlike *Smo*^*bnb/bnb*^ mutant neural tube sections, we observed Olig2 and Nkx6.1 expression in *Inpp5e*^*rdg/rdg*^;*Smo*^*bnb/bnb*^ double mutant embryos (Fig. 5J’, J’’, L’, L’’). Thus, specific ventral cell fates in *Inpp5e*^*rdg/rdg*^ can be specified without Smo. The most parsimonious interpretation of these data is that the absence of Inpp5e function permits derepression of Shh signaling but only to a level permitting an intermediate Shh response.

### Neural tube patterning recovery at E12.5 is dependent on Gli3

Both the *Inpp5e*^*rdg/rdg*^;*Ift172*^*wim/wim*^ and *Inpp5e*^*rdg/rdg*^;*Smo*^*bnb/bnb*^ double mutant embryos express Olig2-positive pMN cells, suggesting that there may be derepression of these cell fates when Inpp5e is lost. In the neural tube, Gli3 is the major repressor of Shh signaling (Litingtung and Chiang, 2000, Persson et al., 2002), and we previously showed that the GliR gradient plays a critical role between E10.5 and E12.5 to properly specify cell fate (Su et al., 2012). We performed Western blots in order to examine Gli processing in *Inpp5e*^*rdg/rdg*^ animals. Using antibodies against Gli2 and Gli3 which detect full-length (185kD Gli2 and 190kD Gli3) and cleaved Gli3 protein (83 kD Gli3R), we evaluated whole embryo protein extracts at E10.5 and E12.5 (Fig. 6A-B). We note that full-length Gli protein does not represent activated Gli protein. We observed no change in full-length Gli2 (Gli2FL) or Gli3 (Gli3FL) between wild type and *Inpp5e*^*rdg/rdg*^ at either E10.5 or E12.5. We saw no change in the 83kD Gli3 band in *Inpp5e*^*rdg/rdg*^ compared to wild type at either E10.5 or E12.5. These data suggest that the overall levels of Gli2FL, Gli3FL and Gli3R are unaltered in homogenized *Inpp5e*^*rdg/rdg*^ embryos.

**Figure 6:**
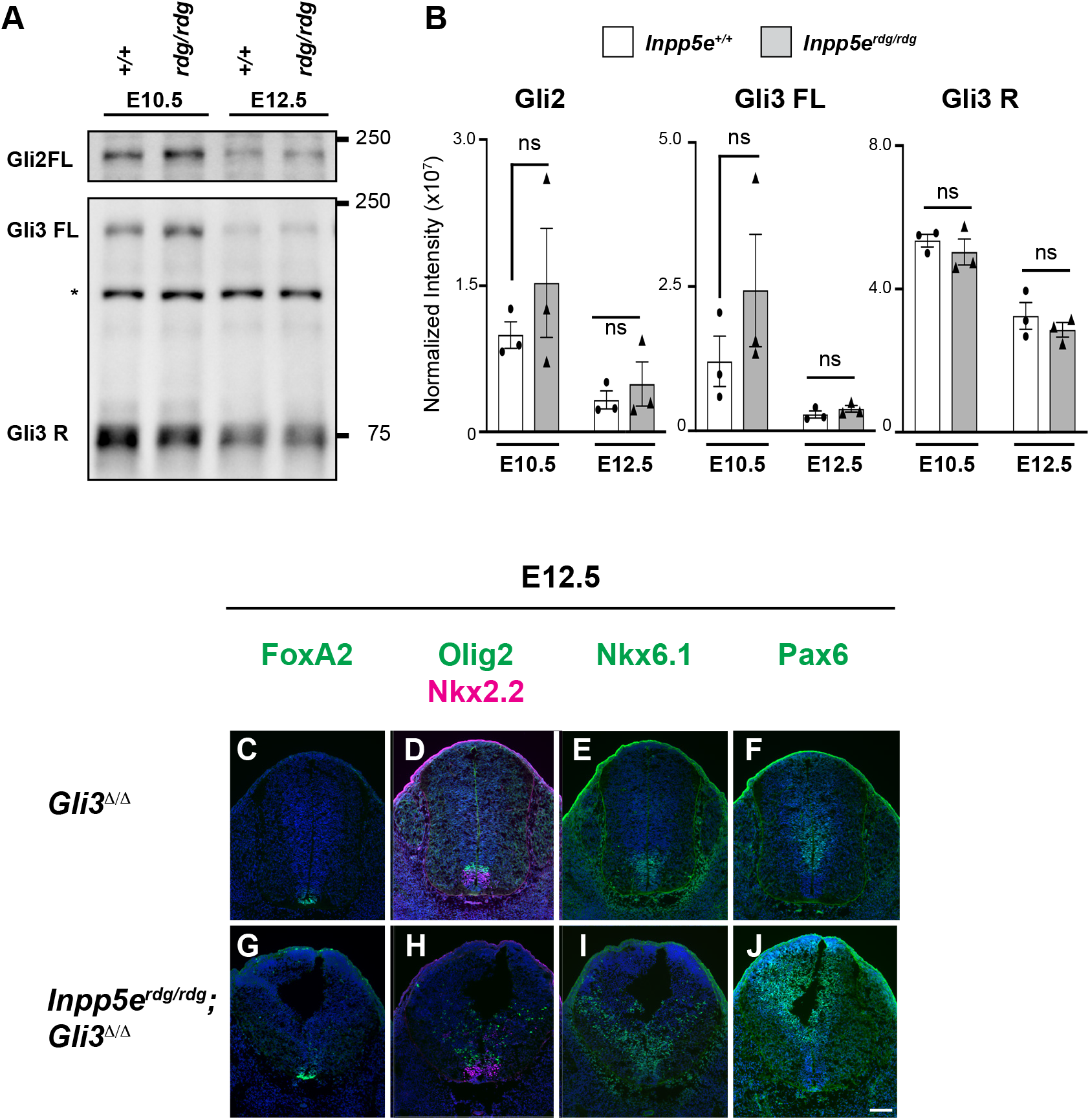
Normal ventral neural patterning in E12.5 *Inpp5e*^*rdg/rdg*^ embryos reflects a recovery of Shh response and is Gli3-dependent. (A) Western blot analysis of Gli2 and Gli3 in wild type and *Inpp5e*^*rdg/rdg*^ whole embryo extracts. Asterisk denotes non-specific band. (B) Quantification of A. Data are mean ± SEM of three biological replicates. ns, not significant. Caudal (hindlimb) sections of E12.5 *Gli3*^*Δ/Δ*^ (C-F) and *Inpp5e*^*rdg/rdg*^;*Gli3*^*Δ/Δ*^ (G-J) neural tubes stained with antibodies against the indicated cell fates. (C-F) FoxA2, Olig2 and Nkx2.2 expression are normal, and Pax6 is expressed correctly in *Gli3*^*Δ/Δ*^ neural tube sections (n=2). (G-I) FoxA2, Olig2 and Nkx6.1 cells are dorsally scattered (expanded) in E12.5 *Inpp5e*^*rdg/rdg*^;*Gli3*^*Δ/Δ*^ neural tube compared to *Gli3*^*Δ/Δ*^ neural tube. The lumen shape of E12.5 *Inpp5e*^*rdg/rdg*^;*Gli3*^*Δ/Δ*^ neural tube resembles that of *Inpp5e*^*rdg/rdg*^ at E10.5 (Fig. 1) (n=3). Scale bar: 100μm.

In order to determine whether Gli3 repressor is functionally altered in *Inpp5e*^*rdg/rdg*^ embryos, we evaluated *Inpp5e*^*rdg/rdg*^;*Gli3*^*Δ/Δ*^ double mutants. At E12.5, we found the *Gli3*^*Δ/Δ*^ mutants specified FoxA2-, Nkx2.2- and Olig2-positive cells in the same domains as the control and *Inpp5e*^*rdg/rdg*^ littermates (Fig. 6C-E, Fig. 3). This is consistent with previous reports of *Gli3*^*Δ/Δ*^ mutants displaying normal specification of these cell fates (Litingtung and Chiang, 2000, Persson et al., 2002). In contrast, we found dorsally scattered FoxA2- and Nkx2.2-positive cells and an expansion of both Olig2- and Nkx6.1-positive cells in E12.5 *Inpp5e*^*rdg/rdg*^;*Gli3*^*Δ/Δ*^ neural tubes (Fig. 6G-I), as well as a dorsal shift in Pax6 staining boundary (Fig. 6J). This indicates the recovery of patterning we observed in *Inpp5e*^*rdg/rdg*^ mutant embryos by E12.5 does not occur in *Inpp5e*^*rdg/rdg*^;*Gli3*^*Δ/Δ*^ mutant embryos. These data demonstrate that the *Inpp5e*^*rdg/rdg*^ recovery phenotype at E12.5 is Gli3-dependent and strongly implicates the GliR gradient in the recovery.

## Discussion

Our data point to Inpp5e regulating the Shh response through a more complicated mechanism than previously appreciated. In vitro, we found *Inpp5e*^*rdg/rdg*^ MEFs did not respond to Shh stimulation, consistent with previous reports that Inpp5e plays a positive role in Shh signal transduction (Garcia-Gonzalo et al., 2015, Dyson et al., 2017, Chavez et al., 2015). However, in vivo we found *Inpp5e*^*rdg/rdg*^ embryos displayed an expansion of ventral cell fates in the neural tube indicating loss of negative regulation of the pathway. Importantly, we observed distinct cell fates requiring high, intermediate and low levels of Shh response in the *Inpp5e*^*rdg/rdg*^ mutant neural tube signifying graded pathway regulation remained. Furthermore, we found that the mispatterning of the E10.5 *Inpp5e*^*rdg/rdg*^ neural tube recovered by E12.5 and that the recovery depended on Gli3, which functions predominantly as a repressor in the neural tube. This finding is consistent with our genetic test of the relationship between Inpp5e and Smo, where we found the *Inpp5e*^*rdg/rdg*^;*Smo*^*bnb/bnb*^ neural tube exhibited cell fates requiring intermediate levels of Shh response such as Olig2 and Nkx6.1. As Smo is essential for Shh signal transduction, this result implies a derepression of intermediate cell fates occurs when Inpp5e function is lost in combination with loss of Smo function. Inpp5e localizes to cilia and we showed that the *Inpp5e*^*rdg/rdg*^ phenotype largely depended on the presence of cilia, consistent with Inpp5e regulating Shh signaling from within the cilium. We showed the *Inpp5e* mutant alleles are sensitive to strain background, which along with the complementation test, enabled us to demonstrate that *Inpp5e*^*rdg*^ is a functional null allele.

The simplest model to explain the expansion of ventral fates we observed in *Inpp5e*^*rdg/rdg*^ mutants is that Inpp5e normally serves as a negative regulator of Shh signal transduction. However, that interpretation is complicated by the normal cell fate specification we observed in E12.5 *Inpp5e*^*rdg/rdg*^ embryos along with our finding that Inpp5e function is required for the Shh response in cell culture which aligns with previously published work (Chavez et al., 2015, Dyson et al., 2017, Garcia-Gonzalo et al., 2015). These data indicate that Inpp5e regulates the Shh response over time and can act as both a positive and negative regulator of the pathway. Thus, any model of Inpp5e function must reconcile these distinct observations.

Other negative regulators of Shh signaling fall into two classes: those whose loss leads to complete constitutive activation of the pathway such as *Ptch1*, *SuFu* or *Gnas* (encoding Gα_s_) mutants, and those whose loss is slightly less severe such as *Tulp3*, *Gpr161* or *Rab23* mutants (Mukhopadhyay et al., 2013, Norman et al., 2009, Patterson et al., 2009, Goodrich et al., 1997, Cooper et al., 2005, Svard et al., 2006). Ptch1, SuFu and Gα_s_ maintain the pathway in an “off” state when ligand is not present so their loss leads to complete pathway activation: increased GliA production, almost no GliR production, and specification of neural fates requiring the highest Shh response. In contrast, Tulp3, Gpr161 and Rab23 adjust the output of the pathway without being essential; they attenuate the pathway. Inpp5e appears to function in this second category: both the *Inpp5e*^*rdg*^ and *Inpp5e*^*ΔEx2-6*^ alleles allow multiple ventral neural cell fates to be specified, indicating that the GliA/GliR ratio is altered consistent with the expanded Shh activity gradient we observed (Dyson et al., 2017). Furthermore, our findings that the recovery of neural patterning in *Inpp5e*^*rdg/rdg*^ embryos between E10.5 and E12.5 is Gli3-dependent along with *Inpp5e*^*rdg/rdg*^;*Smo*^*bnb/bnb*^ mutant embryos exhibiting derepression of Olig2 and Nkx6.1 expression are consistent with lowered GliR production in *Inpp5e*^*rdg/rdg*^ mutants. Reduced GliR production alters the effective GliA/GliR ratio consistent with the expansion of ventral cell fates in *Inpp5e*^*rdg/rdg*^ mutants. The GliA/GliR ratio appears highly variable based on the intermingling of cell fates in the *Inpp5e*^*rdg/rdg*^ neural tube. Taken together, these observations indicate that Inpp5e is an attenuator of Shh signaling in specifying neural cell fates and hint that it is critical for the temporal control of the activator to repressor ratio in the neural tube.

Similar to previous findings in cultured cells, we found *Inpp5e*^*rdg/rdg*^ mutant MEFs exhibited no Shh transcriptional response (Chavez et al., 2015, Dyson et al., 2017, Garcia-Gonzalo et al., 2015). We derived the MEFs from embryos carrying the *Inpp5e*^*rdg*^ allele on an FVB background, the same genetic background on which we observed an expansion of Shh-dependent cell fates in the neural tube. Thus, in contrast to its function in the neural tube, Inpp5e plays a positive role in promoting the Shh response in MEFs. While at the surface contradictory, this adds *Inpp5e*^*rdg*^ to the list of mutations in Shh signal transduction components that reveal distinctions in their neural and fibroblast phenotypes (Gigante et al., 2018, Larkins et al., 2011, Pusapati et al., 2018).

The variance in cell sensitivity to Shh ligand between fibroblasts (NIH/3T3 cells) and neural progenitors lacking Gpr161 is proposed to be due to cell type-specific differences in PKA activity (Pusapati et al., 2018). Differential PKA activity could also explain the distinct phenotypes in fibroblasts and neural progenitors lacking functional Inpp5e. Loss of Inpp5e function in fibroblasts elevates ciliary PIP_2_ levels leading to increased recruitment of Tulp3 and Gpr161, which in turn increases PKA activity resulting in an absence of Shh target gene expression (Garcia-Gonzalo et al., 2015, Mukhopadhyay et al., 2013). While we observed a higher percentage of Tulp3-positive cilia in the *Inpp5e*^*rdg/rdg*^ neural tube compared to wild type, the recruitment of Gpr161 to cilia was limited. Concomitantly, we found a significant increase in the region over which Smo enrichment in cilia was visible in the *Inpp5e*^*rdg/rdg*^ ventral neural tube. The combination of less ciliary Gpr161 and more ciliary Smo would be predicted to result in lower PKA activity in the neural progenitors compared to the MEFs which could render the cells more sensitive to Shh ligand. This would also be consistent with a low level of GliR production in the E10.5 *Inpp5e*^*rdg/rdg*^ neural tube.

Taken together, our data support a model in which loss of Inpp5e function results in the alteration of the normal effective Gli ratio formed from the additive result of the GliA and GliR concentration gradients (Fig. 7). A delay in GliR gradient formation would alter the kinetics of the effective Gli ratio such that there is an initial excess of GliA function that normalizes over time as the standard GliA/GliR ratio forms (Fig. 7B). This model reconciles the seemingly discordant aspects of the Inpp5e phenotypes in several ways. First, in *Inpp5e*^*rdg/rdg*^ mutants at E9.5 and E10.5 low GliR production would derepress known GliR targets: Nkx6.1 and Olig2. This derepression was also evident in the *Inpp5e*^*rdg/rdg*^;*Ift172*^*wim/wim*^ me, low GliR production would increase the effective GliA/GliR ratio which would specify more FoxA2- and Nkx2.2-positive cell fates, as we saw at E9.5 and E10.5 in *Inpp5e*^*rdg/rdg*^ mutants. Second, the fact that the recovery of *Inpp5e*^*rdg/rdg*^ neural patterning is Gli3-dependent argues that the GliR gradient does eventually form. Finally, the fact that *Inpp5e*^*rdg/rdg*^ and *Inpp5e*^*ΔEx7-8/ΔEx7-8*^ MEFs do not respond to Shh ligand is consistent with fibroblasts being more efficient than neural progenitors at GliR production.

**Figure 7:**
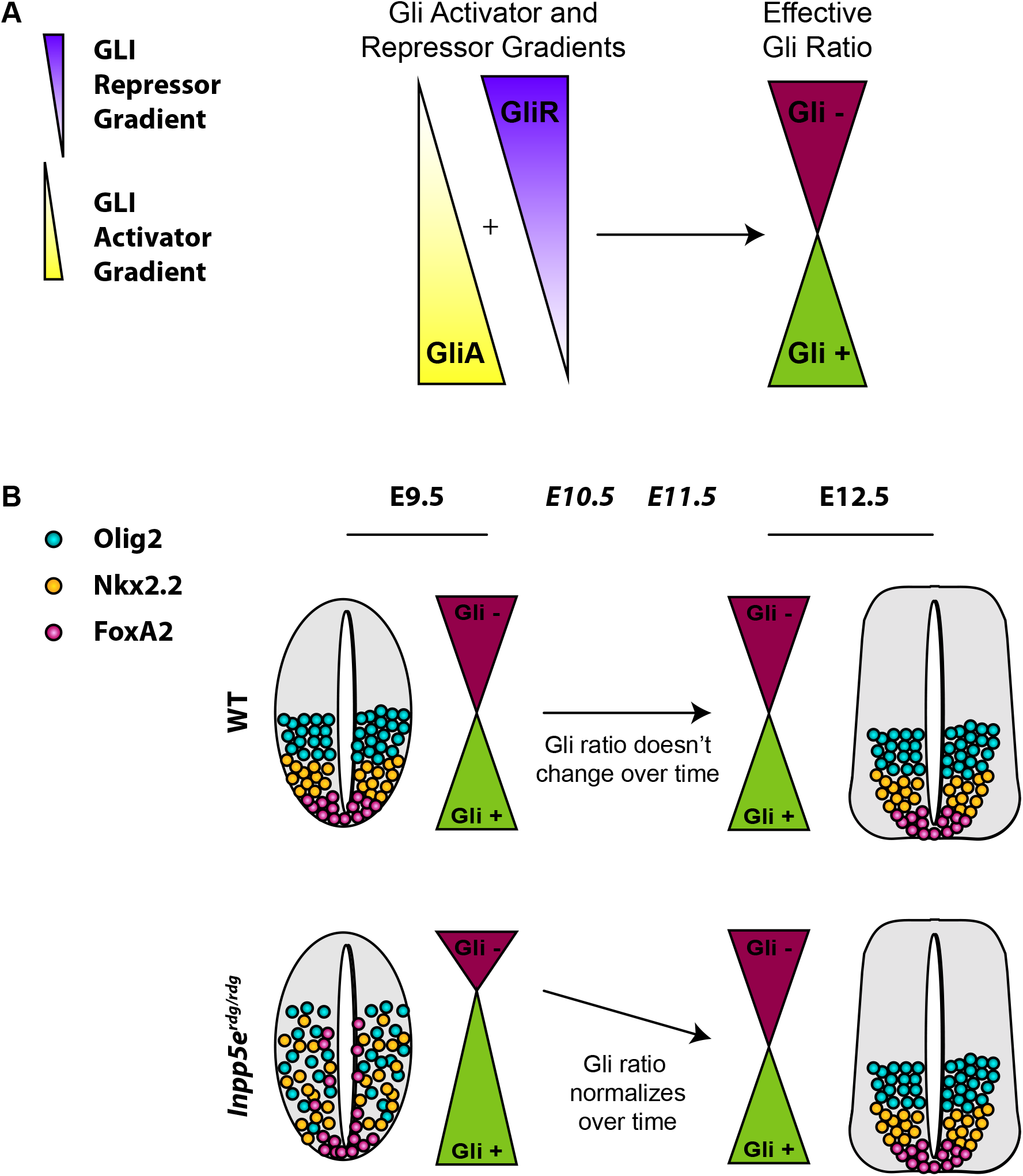
Model for Inpp5e’s role over time in Shh-dependent ventral neural tube patterning. (A) The relative amounts of GliA to GliR specify neural cell fates at any particular position along the ventral-dorsal axis. The combined ratio is the effective Gli ratio (Gli + or Gli -) which integrates the relative ratio of GliA to GliR production. (B) (Top) Normally, the effective Gli ratio doesn’t change over time, leading to normal cell fates. (Bottom) The expanded ventral neural progenitor cell fates in *Inpp5e*^*rdg/rdg*^ mutants at E9.5 normalize by E12.5 due to changes in the effective Gli ratio. While multiple mechanisms are possible, the simplest explanation posits that loss of Inpp5e initially alters GliR gradient production and, over time, the GliR gradient is normalized and the ventral cell fates are returned to their correct positions by E12.5.

The intermingling of cell fates in the *Inpp5e*^*rdg/rdg*^ neural tube suggests that the GliA/GliR ratio is likely quite dynamic and variable from cell to cell. The Gli3-dependent recovery of pattern in E12.5 neural tube argues that Gli3 (the predominant repressor) is biologically significant even if statistically significant differences cannot be seen on Western blots. We note the formal possibility that Inpp5e also changes the kinetics of GliA production; however, changes in the kinetics of GliR production are sufficient to explain the apparent increase in the levels of effective GliA resulting in ventral cell fate expansion. Cells can also integrate the level of Shh signaling over time by taking into account the duration of Gli activity in a process known as temporal adaptation (Stamataki et al., 2005, Dessaud et al., 2007). Cells exposed to lower concentrations of Shh lose Gli responsiveness faster than cells exposed to high concentrations of Shh, allowing cells to interpret both concentration and duration of exposure (Dessaud et al., 2007). Thus, the model we propose awaits techniques that monitor GliA and GliR in situ at the cellular level and over time to tease these two possibilities apart.

Our model can reconcile some additional discrepancies. Unlike the general expansion of neural cell fates we saw in the *Inpp5e*^*rdg/rdg*^ mutants, the *Inpp5e*^*ΔEx2-6/ΔEx2-6*^ mutant specifies no floor plate (requiring the highest levels of Shh signaling) and an expansion of more intermediate cell fates (Dyson et al., 2017). That allele, like the *Inpp5e*^*ΔEx7-8*^ allele, is on a C57BL/6 genetic background and both display perinatal lethality (Dyson et al., 2017, Jacoby et al., 2009). We found the *Inpp5e*^*rdg/rdg*^ and *Inpp5e*^*ΔEx7-8/ΔEx7-8*^ embryos both died at E13.5 on an FVB background but viable homozygous animals were present at E16.5 on a C57BL/6 background. In the context of our model, these data would predict genetic background alters the kinetics of GliA/GliR ratio production. This is consistent with Inpp5e acting as an attenuator of the Shh pathway.

The relationship between Inpp5e and Smo is both straightforward and enigmatic, highlighting the complexity of Inpp5e function. This may be best illustrated by the fact that we found slightly less than half as many *Inpp5e*^*rdg/rdg*^;*Smo*^*bnb/bnb*^ double mutant embryos at E9.5 as expected. These missing embryos are likely biologically significant and further work is needed to define their time of lethality. Nevertheless, their loss is consistent with Inpp5e playing an important role in relation to time. Previous work showed that many phenotypes exhibited by embryos expressing constitutively activated Smo (*SmoM2* allele) can be rescued by *Inpp5e*^*ΔEx2-6/ΔEx2-6*^, consistent with our finding that the *Inpp5e*^*rdg/rdg*^ phenotype is Smo-dependent and arguing that Inpp5e functions at a step downstream of Smo (Dyson et al., 2017). It also suggests that Inpp5e loss reduces the activator output of the pathway providing in vivo evidence that Inpp5e plays a positive role in regulating Shh signal transduction. However, the expansion of ventral neural fates displayed by E10.5 *Inpp5e*^*ΔEx2-/6/ΔEx2-6*^ or *Inpp5e*^*rdg/rdg*^ mutants implies Inpp5e negatively regulates the Shh response. This is likely explained, in part, by the links between PI(4)P, Ptch1 and Smo along with the diminished ciliary PI(4)P in the absence of Inpp5e function (Garcia-Gonzalo et al., 2015, Chavez et al., 2015, Yavari et al., 2010, Jiang et al., 2016). Such roles occur upstream of activated Smo so would be masked in the context of the *SmoM2* allele.

The increase in ciliary Smo in *Inpp5e*^*rdg/rdg*^ neural tubes is consistent with the expansion of ventral cell fates. Ciliary Smo enrichment normally clears Gpr161 from cilia, thereby removing the pathway’s basal repression machinery. However, we observed 34% of cilia were Gpr161-positive in *Inpp5e*^*rdg/rdg*^. Inpp5e knockdown in IMCD3 cells leads to the simultaneous presence of both Smo and Gpr161 in cilia (Badgandi et al., 2017). Furthermore, ciliary clearance of Gpr161 can be uncoupled from Shh activation (Pusapati et al., 2018). These data together suggest that loss of Inpp5e uncouples the normal regulatory mechanism between Smo and Gpr161 and could explain how signaling occurs in the presence of ciliary Gpr161 (Badgandi et al., 2017, Pal et al., 2016). Our finding of Tulp3 in almost all *Inpp5e*^*rdg/rdg*^ neural tube cilia raises another possibility. Tulp3 is known to traffic multiple GPCRs into the cilium as well as other molecules known to regulate the Shh pathway, so perhaps Tulp3 is regulating another modulator of the Shh response (Mukhopadhyay et al., 2010, Hwang et al., 2019, Legue and Liem, 2019, Han et al., 2019, Badgandi et al., 2017). Taken together, this would explain the paradoxical increase in Shh signaling in the presence of high Tulp3 and Gpr161 seen in our embryos.

Inpp5e functions within cilia to maintain the PI(4)P membrane composition and our analysis is consistent with Inpp5e functioning within cilia to regulate Shh signaling. Other enzymes that impact PI(4)P are known to regulate Hedgehog (Hh) in Drosophila, where the pathway does not rely on cilia, suggesting that PI signaling is an ancient mechanism for Hh regulation (Yavari et al., 2010). The fact that our data argue Inpp5e plays a critical role for Shh signal transduction over time is provocative. Other than the reliance on cilia, the fundamental distinction between vertebrate and invertebrate Hh signaling is that vertebrates use Hh for long-range signaling. Previous data showed that changing the kinetics of ciliary traffic can alter the output of the Shh pathway (Ocbina et al., 2011). Thus, we speculate that vertebrate Hh signaling through/via cilia may enable cells to control the duration or timing of the Shh signal.

In conclusion, our data provide genetic evidence that Inpp5e attenuates Shh signaling in the developing mouse neural tube. Our data add to the existing function of Inpp5e and argue that it plays both positive and negative regulatory roles in Shh signal transduction, likely through controlling the timing of Gli processing and thus the sensitivity of cells to respond to Shh ligand, both in the dorsal-ventral axis and over time. These data also highlight the phenotypic variability among Inpp5e mutant cell types and alleles, expanding our knowledge on the role of Shh signaling duration in regulating ventral neural cell fate.

## Acknowledgments

We are grateful to K. Anderson for the Smo antibody, S. Mukhopadhyay for the Gpr161 antibody and J. Eggenschwiler for the Tulp3 antibody. We are also grateful to S. Mukhopadhyay for critical comments on the manuscript along with members of the Caspary lab. We thank C. Lin for his initial *Inpp5e*^*rdg*^ observations. The content is solely the responsibility of the authors and does not necessarily reflect the official views of the National Institute of Health.

---

**Supplemental Figure 1:**

***Inpp5e*^*ΔEx7-8/ΔEx7-8*^ embryos show expanded ventral cell fates at E10.5 and recovery at E12.5.**

Caudal (hindlimb) sections of E10.5 and E12.5 neural tube sections stained with antibodies against indicated cell fates. (A-J) E10.5 *Inpp5e*^*ΔEx7-8/ΔEx7-8*^ mutants show expanded ventral cell fates and Shh activity compared to wild type embryos (wild type n=6, *Inpp5e*^*ΔEx7-8/ΔEx7-8*^ n=4). This phenotype is similar to what we observed in *Inpp5e*^*rdg/rdg*^ mutants (see Fig. 1F-J). (K-T) At E12.5, these expanded cell fates have returned to normal, with few cells scattered dorsally (wild type n=6, *Inpp5e*^*ΔEx7-8/ΔEx7-8*^ n=2). Scale bars: 100μm.

## Materials and Methods

### Mouse lines and maintenance

All mice were cared for in accordance with NIH guidelines and Emory’s Institutional Animal Care and Use Committee (IACUC). Alleles used were: *Inpp5e*^*rdg*^ [MGI:6295836], *Ptch1*^*LacZ*^ (*Ptch1*^*tm1Mps*^) [MGI:1857447], *Inpp5*^*eΔEx7-8*^ (*Inpp5e*^*tm1.2Ssch*^) [MGI: 4360187], *Ift172*^*wim*^ [MGI: 2682066], *Smo*^*bnb*^ [MGI: 2137553], *Gli3*^*fl*^ (*Gli3*^*tm1Alj*^) [MGI: 3798847] and *CAGGCre-ER*^*TM*^(Tg(CAG-cre/Esr1&)5Amc) [MGI: 2182767]. *Inpp5e*^*ΔEx7-8/+*^ animals were generated by crossing *Inpp5e*^*fl/fl*^ (*Inpp5e*^*tm1.1Ssch*^) [MGI: 4360186] animals to *CAGGCre-ER*^*TM*^ animals and treating pregnant dams with tamoxifen as previously described (Su et al., 2012). *Inpp5e*^*fl/fl*^ animals were received at Emory University and rederived on C57BL/6J. They, and the derived *Inpp5e*^*ΔEx7-8/+*^ mice, are maintained on FVB/NJ. *Inpp5e*^*rdg/rdg*^;*Gli3*^*Δ/Δ*^ embryos were generated by crossing *Inpp5e*^*rdg/+*^*;Gli3*^*fl/+*^ and *Inpp5e*^*rdg/+*^*;Gli3*^*fl/+*^;*CAGGCre-ER*^*TM*^ animals and treating pregnant dams with tamoxifen at E7.5 as previously described (Su et al., 2012). *CAGGCre-ER*^*TM*^, *Ptch1*^*LacZ*^, *Ift172*^*wim*^, *Smo*^*bnb*^, and *Gli3*^*fl*^ were on a C3H/HeJ background when this project began and are currently maintained with *Inpp5e*^*rdg*^ on FVB/NJ. Genotyping was performed as previously described or with Transnetyx, Inc. (Goodrich et al., 1997, Blaess et al., 2008, Jacoby et al., 2009, Kasarskis et al., 1998, Hayashi and McMahon, 2002, Sun et al., 2012). Timed mating of heterozygous intercrosses was performed with animals less than a year old to generate embryos of the indicated embryonic stage.

### Mouse dissection, X-gal staining and immunofluorescence

Embryos were dissected in cold phosphate-buffered saline (PBS) and processed for either X-gal staining or immunofluorescence.

Embryos were stained with X-gal as previously described (Goodrich et al., 1997). After fixing in 4% paraformaldehyde (PFA) overnight at 4°C, embryos were incubated in 30% sucrose in 0.1M phosphate buffer, pH 7.3, at 4°C overnight prior to being embedded in OCT (Tissue-Tek) and 40μm sections were obtained on a Leica CM1850 cryostat.

For immunofluorescence, embryos were fixed for 1 h in 4% PFA on ice. Embryos were processed through sucrose and OCT as above, before sectioning at 10μm. Sections were incubated with primary and secondary antibodies diluted in PBS with 0.1% Triton X and either 1% or 10% heat inactivated goat serum. The following primary antibodies were used: mouse anti-Shh clone 5E1 (1:10), mouse anti-Nkx2.2 clone 74.5A5, mouse anti-Nkx6.1 clone F65A2, and mouse anti-Pax6 (1:100) from Developmental Studies Hybridoma Bank; rabbit anti-FoxA2 (Cell Signaling, #3143; 1:500), rabbit anti-Olig2 (Millipore, AB9610; 1:300), rabbit anti-Smo (kind gift from K. Anderson; 1:1000), mouse anti-Arl13b (Neuromab, N295B/66; 1:2000), rabbit anti-Tulp3 (kind gift from J. Eggenschwiler; 1:500), and rabbit anti-Gpr161 (kind gift from S. Mukhopadhyay; 1:200). The secondary antibodies were conjugated to Alexa Fluor 488, 568, or 594 (ThermoFisher; 1:200). Hoechst 33342 (Sigma; 1:3000) was included in the incubation of slides with secondary antibody.

### Smo, Gpr161 and Tulp3 quantification

Three sections from each embryo (wild type n=3, *Inpp5e*^*rdg/rdg*^ n=3) were used for quantification of ciliary Smo, Tulp3 and Gpr161. To quantify the Smo expansion, the measurement tool in FIJI image processing software package was used to draw a line around the entire neural tube lumen (Schindelin et al., 2012). The distance over which ciliary Smo enrichment was observed was determined and reported as a percentage of the total distance of the neural tube lumen. To quantify Tulp3 and Gpr161, the number of Arl13b positive luminal cilia which were also positive for either Tulp3 or Gpr161 in the ventral 10% of the neural tube were counted using the counting plugin in FIJI. Three sections from each embryo (wild type n=3, *Inpp5e*^*rdg/rdg*^ n=3), between 400 and 500 cilia, were counted for each condition per genotype. Statistical significance was evaluated on three biological replicates in PRISM v8.1.1 using two tailed unpaired t-test with Welch’s correction.

### MEF generation, RNA isolation and qPCR quantitation

Mouse embryonic fibroblasts (MEFs) were isolated from E12.5 embryos and immortalized as previously described (Mariani et al., 2016). Genotypes were verified by PCR. For Shh treatment, *Inpp5e*^*rdg/+*^ and *Inpp5e*^*rdg/rdg*^ MEFs were grown at a density of 0.5 × 10^6^ cells/mL and treated for 24 h with Shh-conditioned medium containing 0.5% fetal bovine serum (Larkins et al., 2011).

For qPCR, whole RNA was extracted from MEFs and qPCR was carried out as previously described (Bay et al., 2018, Gigante et al., 2018). The following primers were used (5′-3′): Gli1 (GCCACACAAGTGCACGTTTG and AAGGTGCGTCTTGAGGTTTTCA); Gapdh (CGTCCCGTAGACAAAATGGT and GAATTTGCCGTGAGTGGAGT) (Bay et al., 2018). Each reaction was performed in technical triplicate. Gli1 values were normalized to Gapdh within each sample. Statistical significance was evaluated in PRISM v8.1.1 by applying a two-way ANOVA with Tukey correction for multiple analysis on three biological replicates.

### Western Blotting

Western blotting was performed as previously described (Chang et al., 2016, Mariani et al., 2016, Bay et al., 2018) with the following antibodies: Gli2 (R&D Systems, AF3635, 1:500), Gli3 (R&D Systems, AF3690, 1:1000), and HRP-conjugated donkey anti-goat IgG (Jackson ImmunoResearch, 1:5000). Lysates were made using RIPA buffer with Roche protease inhibitors (Chang et al., 2016). Values displayed are volume intensity as measured from a chemiluminescent image and normalized to total protein as measured on a stain-free gel. Statistical significance was evaluated in PRISM v8.1.1 by applying a two-way ANOVA with Tukey correction for multiple analysis on three biological replicates.

